# Morphine reprograms brain-derived extracellular vesicles from cargo linked to synaptic remodelling in the prefrontal cortex

**DOI:** 10.1101/2025.09.12.675684

**Authors:** Pedro Henrique Gobira, Savio Bastos, Rachele Rossi, Christian Bjerggaard Vægter, Fenghua Chen, Nicole Rodrigues da Silva, Heidi Kaastrup Müller, Samia Joca, Thomas Boesen, Thiago Wendt Viola, Yan Yan, Martin R. Larsen, Victor Corasolla Carregari, Rodrigo Grassi-Oliveira

**Affiliations:** Translational Neuropsychiatry Unit, Department of Clinical Medicine, Aarhus University, Aarhus, Denmark; Aarhus University, Biomedicine, Aarhus, Denmark; Interdisciplinary Nanoscience Center -iNANO, Aarhus University, Aarhus; Nanopsychiatry Lab, School of Medicine, Pontifical Catholic University of Rio Grande do Sul (PUCRS), Brazil; EVGenomics ApS, Aarhus, Denmark; Department of Biochemistry and Molecular Biology, University of Southern Denmark, Odense, Denmark

**Keywords:** extracellular vesicles, morphine, prefrontal cortex, transcriptomics, proteomics, synaptic plasticity

## Abstract

Extracellular vesicles (EVs) released by neurons and glial cells mediate intercellular communication in the brain and regulate synaptic function, neuronal survival, and neuropathological processes. Although chronic opioid exposure induces widespread neuroadaptations, the contribution of brain-derived EVs (BDEVs) to these processes remains largely unknown. Here, we isolated BDEVs from the prefrontal cortex of rats chronically exposed to morphine and performed integrative transcriptomic and proteomic analyses of their molecular cargo. Total RNA sequencing combined with unbiased proteomics revealed that morphine profoundly reprograms the BDEV transcriptome and proteome, enriching pathways related to synaptic plasticity, endoplasmic reticulum (ER) stress, mitochondrial dysfunction, and neurodegeneration. Among the most prominent alterations, the synaptic regulator *ARC* was consistently modulated at the mRNA level, while the ER stress marker *HSPA5* was altered at both mRNA and protein levels. Functional assays further demonstrated that BDEVs derived from morphine-treated rats were sufficient to reconfigure transcriptional programs in naïve cortical neurons, affecting genes associated with synaptic remodeling and excitability. Collectively, these findings provide the first evidence that chronic opioid exposure reprograms BDEV cargo in a brain region critical for addiction and that these vesicles can propagate transcriptional reorganization to recipient neurons. BDEVs thus emerge as active mediators of morphine-induced neuroadaptations and as potential targets for biomarker discovery and therapeutic intervention in opioid use disorder.

## Introduction

Opioid use disorder (OUD) remains a major public health crisis, driven by persistent neuroadaptations that underlie compulsive drug seeking and relapse [1,2]. Although the synaptic and receptor-level effects of opioids have been extensively characterized [3,4] , the intercellular mechanisms through which prolonged exposure remodels brain circuits are still not fully understood. Bridging this gap requires examining how cells communicate beyond classical synaptic signaling, through molecular messengers capable of propagating neuroadaptations across neural networks.

Extracellular vesicles (EVs) are nanosized particles secreted by virtually all cell types, including neurons and glia, that transport proteins, RNAs, and lipids between cells [5]. Within the central nervous system, brain-derived EVs (BDEVs) regulate synaptic plasticity, stress responses, and neuronal survival [6–8]. Given their ability to mediate intercellular communication, BDEVs have emerged as key candidates for understanding how chronic drug exposure reshapes brain function. Disruptions in these processes have been increasingly implicated in psychiatric conditions, including substance use disorders [8]. However, although emerging evidence suggests that drugs of abuse can modify the molecular cargo of EVs, the link between such changes and broader neurobiological processes remains poorly defined [9].

Most prior work has focused on small RNAs, particularly microRNAs, based on the assumption that EVs are selectively enriched for these species [10,11]. Yet this focus may overlook critical layers of regulation, as recent studies demonstrate that BDEVs can also deliver intact mRNAs, such as the synaptic regulator ARC, which are translated in recipient neurons and contribute to neuroadaptive remodelling across interconnected brain regions [12,13]. These findings suggest that EVs may act as functional conveyors of complex molecular programs, extending opioid-induced neuroplasticity beyond synaptic boundaries.

Here, we test the hypothesis that chronic morphine exposure reprograms the transcriptomic and proteomic composition of BDEVs and that these vesicles are sufficient to drive transcriptional changes in recipient neurons. We isolated BDEVs from the prefrontal cortex (PFC), a region central to addiction-related plasticity, and conducted integrative RNA sequencing and proteomic profiling. Functional validation was performed by exposing primary cortical neurons to BDEVs derived from morphine- or vehicle-treated rats, followed by transcriptional analyses. Together, these experiments identify BDEVs as potential mediators of opioid-induced neuroadaptations, providing novel insights into the pathophysiology of OUD and highlighting opportunities for biomarker development and therapeutic innovation.

## Methods and Materials

Methods are summarized below. A complete description of all experimental procedures is available in Supplementary Materials.

### Animals and drug treatment

Male Sprague–Dawley rats (8 weeks old, Janvier Labs) were pair-housed under standard conditions with ad libitum access to food and water. After two weeks of habituation, animals received intraperitoneal injections of saline or escalating doses of morphine (10–50 mg/kg, twice daily for 5 days). Four hours after the final injection, rats were sacrificed, and bilateral PFC was dissected and frozen at −80 °C. All procedures were approved by the Danish Animal Experiments Inspectorate and complied with EU Directive 2010/63/EU.

### Isolation and characterization of brain-derived extracellular vesicles (BDEVs)

BDEVs were isolated from PFC tissue by enzymatic dissociation, differential centrifugation, filtration, and ultracentrifugation, as previously described [14]. Vesicle purity and morphology were assessed by BCA protein assay, Western blotting, transmission electron microscopy (TEM), and nanoparticle tracking analysis (NTA). Full protocols are provided in Supplementary Methods.

### RNA extraction, sequencing, and analysis

RNA was extracted from BDEVs, DNA was removed, and total RNA libraries were prepared using a stranded RNA-seq kit. Libraries were sequenced on an Illumina NovaSeq X Plus platform (150 bp, paired-end). Quality control, differential gene expression analysis (DESeq2), and pathway enrichment (Metascape, GO, and Reactome) followed standard procedures, with significance defined at FDR-adjusted p < 0.1 and log2FC ±0.5.

### Proteomic analysis

BDEV proteins were extracted, digested with trypsin, and analyzed by LC–MS/MS on an Orbitrap Q Exactive HF system. Data were searched against the Rattus norvegicus UniProt database and filtered at 1% FDR. Proteins were quantified based on unique peptides, and enrichment analyses were conducted using Cytoscape, IPA, and R-based workflows.

### Primary neuronal cultures and BDEV treatment

Cortical neurons were prepared from embryonic day 17 (E17) C57BL/6 embryos and plated in Neurobasal medium with B27 supplement. On DIV7, neurons were treated with BDEVs isolated from saline- or morphine-exposed rats, and RNA was extracted 24 h later.

### NanoString gene expression profiling

Gene expression in primary neurons was measured using the NanoString nCounter® Mouse Neuropathology Panel (770 genes). Data were normalized to housekeeping genes using nSolver™ 4.0, and pathway enrichment was conducted in Reactome.

## Results

### Characterization of EVs from the prefrontal cortex of chronically morphine-treated rats

BDEVs were isolated from the PFC of rats via ultracentrifugation as previously described [14]. We used NTA to determine the size distribution and particle number, with the latter normalized to the individual PFC weight. Results revealed no significant differences between animals treated with morphine and those treated with vehicle **(Fig. 1B).** Using TEM, we characterized the BDEVs, revealing their characteristic morphology (**Fig. 1C**). We also analyzed BDEVs via specific positive and negative markers via Western blotting. As expected, the positive markers flotillin-1 and Hsp70 were detected in the BDEV fractions, whereas the negative marker GM130 was absent from the BDEV fractions but present in the whole cortex lysates (**Fig. 1D**). Together, these findings confirm the reliability of our isolation technique.

**Figure 1.**
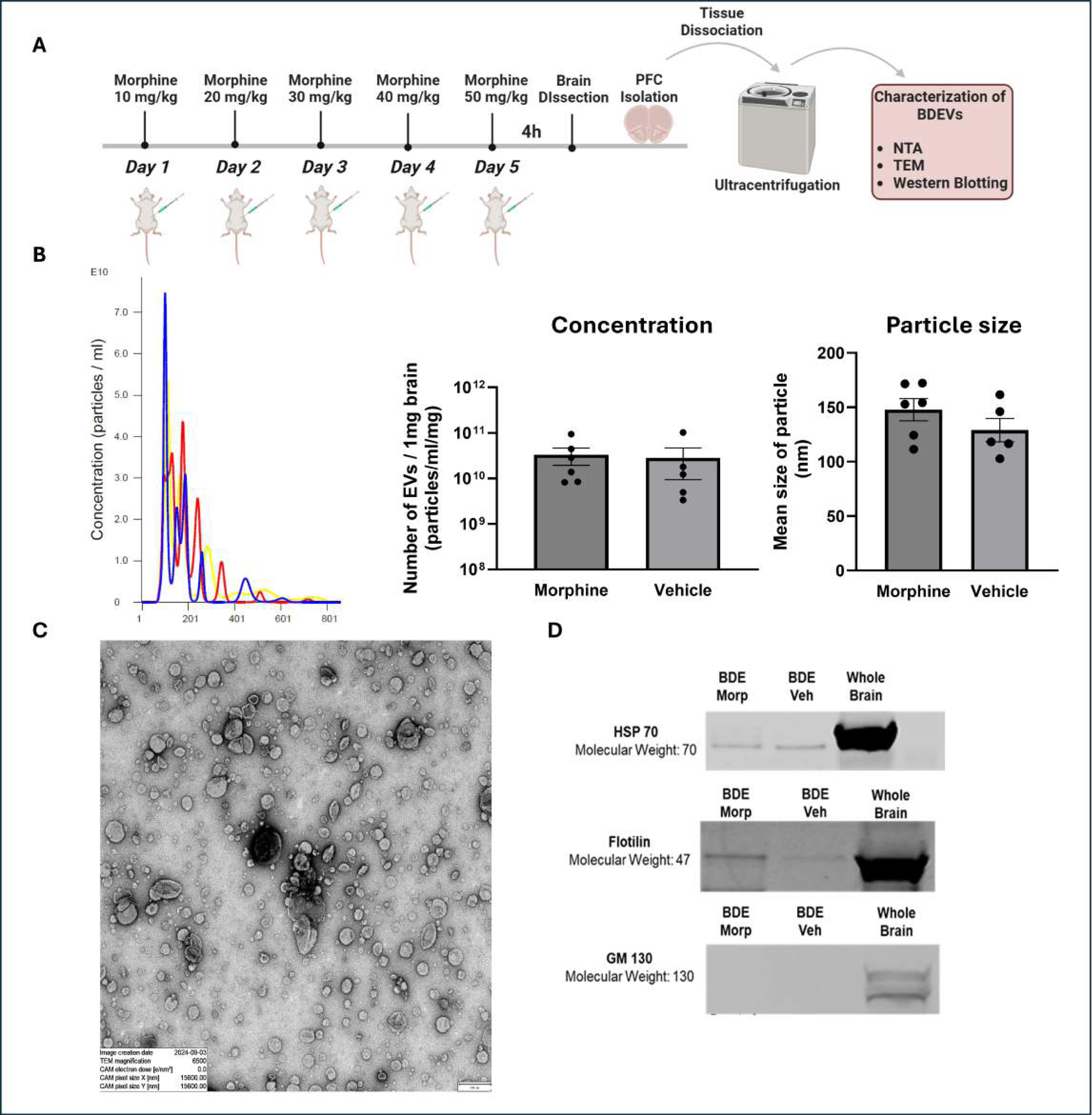
Characterization of BDEVs from rat prefrontal cortex. (A) Experimental design. (B) Nanoparticle tracking analysis (NTA) showing no significant changes in concentration or size distribution of isolated BDEVs between groups.(C) Representative electron micrographs of EVs. (D) Western blot analysis confirming the presence of EV markers Flotillin and HSP70, and absence of the exclusion marker GM130 in the EV samples.

### Effect of morphine treatment on the RNA profile of BDEVs

The RNA-seq analysis identified a total of 12,224 RNA molecules that surpassed the quality control threshold for downstream analysis, comprising 49 long noncoding RNAs (0.40%), 94 small noncoding RNAs (0.77%), 11,948 protein-coding mRNA molecules (97.74%), and other RNAs (e.g., ribosomal RNA, transfer RNA, mitochondrial RNA, and pseudogenes—1.09%). Prior to conducting the differential gene expression (DGE) analysis, the RNA-seq data were deconvoluted from publicly available single-cell RNA sequencing data from rodent brains. We analyzed the proportions of microglia, glutamatergic corticothalamic (CT), intratelencephalic (IT), and pyramidal tract (PT) neurons, as well as GABAergic interneurons. The proportions of oligodendrocytes, astrocytes, and glutamatergic NP neurons were negligible. No significant group differences in cell-type proportions were observed (all p values > 0.05; see **Fig.** 2A). However, we observed that almost 40% of the RNA-seq data were from BDEVs originating from GABAergic interneurons.

**Figure 2.**
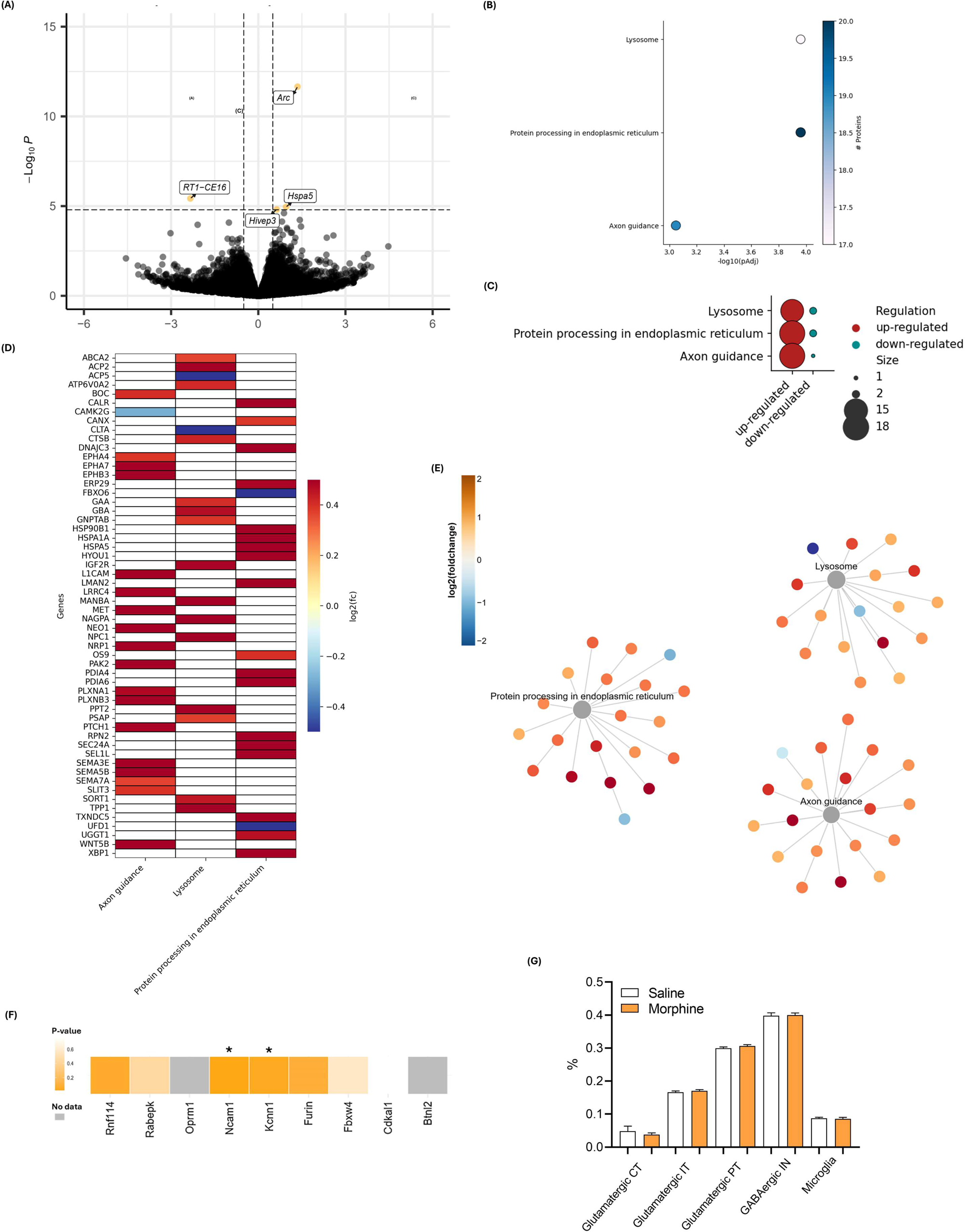
**Effect of morphine treatment on the RNA profile of BDEVs** (A) Volcano plot of differentially expressed mRNAs in BDEVs from morphine-treated versus control animals. (B) KEGG pathway enrichment analysis of morphine-induced RNA alterations. (C) Dot plot showing upregulated and downregulated transcripts within enriched pathways. (D) Heatmap of significantly dysregulated mRNAs across enriched pathways (log₂ fold-change). (E) Network visualization of enriched pathways and associated transcripts. (F) Integration with a human GWAS for opioid use disorder highlights risk-associated genes differentially expressed in BDEVs. (G) RNA deconvolution estimates brain cell-type composition, with nearly 40% of transcripts inferred from GABAergic interneuron-derived BDEVs.

A comprehensive analysis revealed 625 RNA transcripts with unadjusted p values less than 0.05 (Supplemental Table 1), indicating an effect of morphine treatment on the EV transcriptomic cargo. Four hits were identified by DGE analysis as having significant group differences that surpassed the FDR correction thresholds: Arc, Hspa5, Hivep3, and RT1-CE16 **(Fig. 2B)**. These four molecules are mRNAs derived from protein-coding genes. Arc, Hspa5, and Hivep3 were upregulated in EVs from morphine-treated animals, whereas RT1-CE16 was downregulated. Additionally, we cross-referenced our RNA-seq data with a large-scale human genome-wide association study (GWAS) meta-analysis of opioid use disorder [15] to identify differentially regulated transcripts originating from previously identified genes associated with the disorder, such as Oprm1, Cdkal1, Btnl2, Rabepk, Fbxw4, Ncam1, Furin, Kcnn1, and Rnf114. Among these, two transcripts, Ncam1 (log2FC = 0.45; p value = 0.02) and Kcnn1 (log2FC = 0.49; p value = 0.04), had unadjusted p values below 0.05, indicating a greater abundance of EVs in morphine-treated animals than in control animals **(Fig. 2C).**

Pathway enrichment analysis was conducted using all RNAs with an unadjusted p value < 0.05 on the basis of the KEGG 2019 mouse database (*Rattus norvegicus* gene codes were converted to mice before enrichment). Among the enriched pathways, only Lysosome, Protein Processing in the Endoplasmic Reticulum, and Axon Guidance were significantly represented, predominantly comprising upregulated transcripts (Figure y1 a-b). Notably, the enriched RNAs within each pathway were mutually exclusive, with no individual RNA shared across multiple pathways. This suggests a limited overlap and low interpathway correlation among the transcriptional alterations observed (Figure y2a-b).

### Effect of morphine treatment on the protein profile of BDEVs

A total of 1007 proteins were reliably quantified, 379 of which passed the quality control criteria for downstream analysis. Among these, 82 proteins were found to be dysregulated on the basis of unadjusted p values < 0.05 (Supplemental Table 2), including 44 downregulated and 38 upregulated proteins. However, none of these changes remained statistically significant after false discovery rate (FDR) correction. Consequently, we adopted a more permissive threshold, selecting proteins with unadjusted p values < 0.01 and an absolute log2-fold change (log2FC) ≥ 0.5 to highlight potentially relevant alterations, which is consistent with prior proteomic studies applying exploratory thresholds in complex biological models [16,17]. Using these criteria, ten proteins were identified as differentially regulated in BDEVs following morphine exposure (Fig. 3). Among these proteins, Krt33a, Fabp5, Krt72, Sprr1a, and Pkp1 were downregulated, whereas Stx1b, Ywhag, Grm8, Hspa5, and Pdia3 were upregulated.

**Figure 3.**
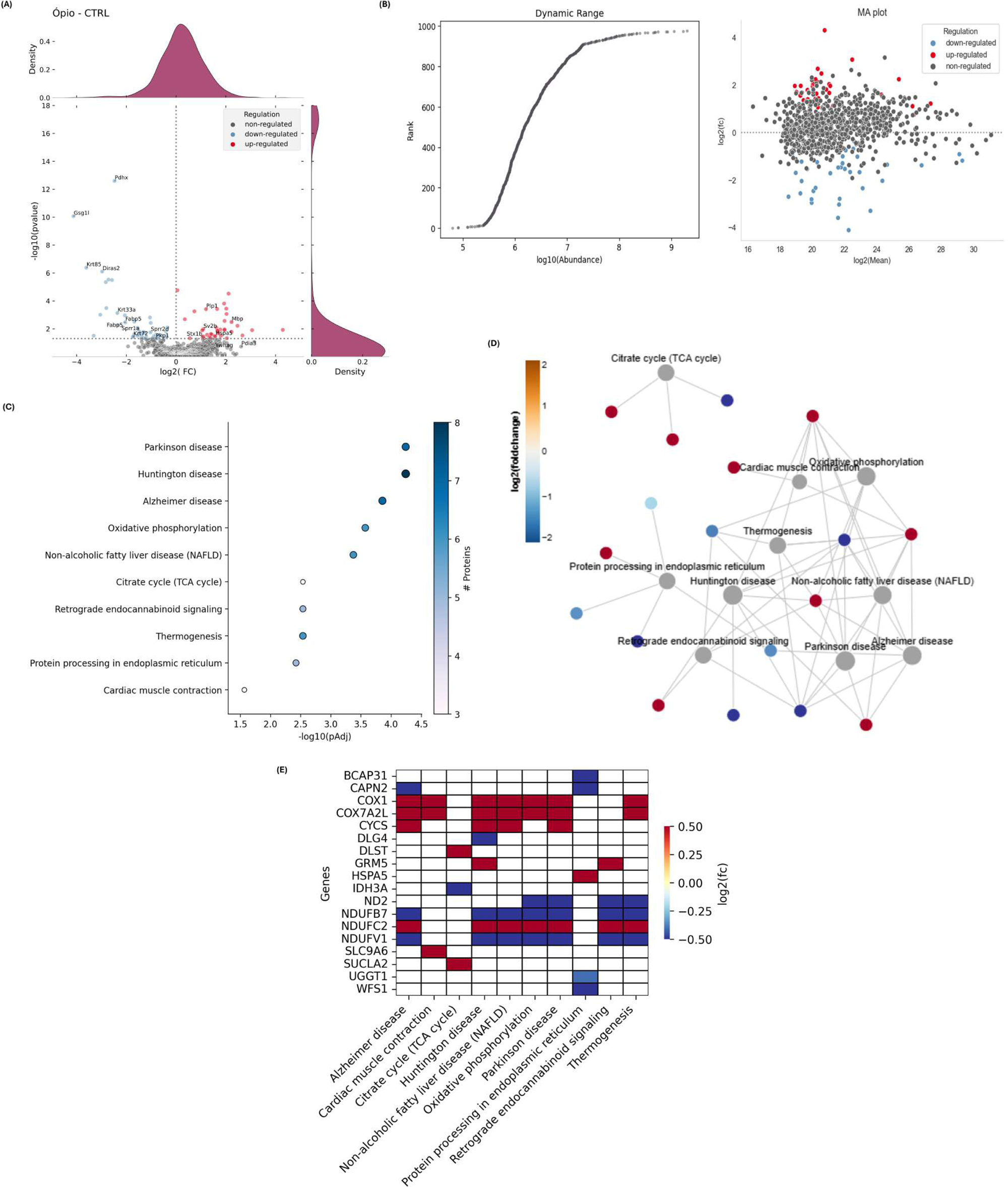
**Effect of morphine treatment on the protein profile of BDEVs** (A) Volcano plot showing differentially expressed proteins in BDEVs from morphine-treated versus control animals. (B) Dynamic range of protein abundance (left) and MA plot (right), highlighting significantly dysregulated proteins. (C) KEGG pathway enrichment analysis identifying pathways associated with morphine-induced protein alterations. (D) Network visualization of enriched pathways and associated proteins. Gray nodes represent KEGG pathways; colored nodes represent proteins, with node color indicating log₂ fold-change (red = upregulated, blue = downregulated). (E) Heatmap of log₂ fold-change values for dysregulated proteins across enriched KEGG pathways.

A volcano plot was used to visualize the overall distribution of protein abundance changes, clearly highlighting the most significantly dysregulated proteins (p value < 0.05). Furthermore, dynamic range analysis revealed a broad spectrum of protein abundances, indicating that both low- and high-abundance proteins were successfully identified and quantified, an important aspect of exosome proteomics that enables the detection of diverse cargo molecules (Figure 3a-b) [18,19].

To elucidate the biological implications of these morphine-associated protein alterations, pathway enrichment analysis was performed via the KEGG Mouse 2019 database [20]. The dysregulated proteins were significantly associated with pathways involved in neurodegenerative diseases, oxidative phosphorylation, nonalcoholic fatty liver disease, the citrate (TCA) cycle, retrograde endocannabinoid signaling, thermogenesis, and protein processing in the endoplasmic reticulum (Figure 3 C).

Inspection of the dysregulation patterns revealed notable enrichment of mitochondrial proteins, particularly members of the NDUF family, as well as COX1 and COX7A2L (Figure 3 D-E), which play essential roles in oxidative phosphorylation and energy metabolism, mechanisms implicated in neuropsychiatric disorders [21–23]. Additionally, the upregulation of the GRM5-encoded metabotropic glutamate receptor in the exosomal cargo of morphine-exposed subjects suggests potential activation of the phosphatidylinositol–calcium second messenger signaling pathway, which is known to influence synaptic plasticity and drug-related neuroadaptation [24,25].

### Integrative analysis of differentially regulated RNAs and proteins

Proteomic analysis identified 1,084 proteins, of which 1,007 were quantifiable. Notably, more than 85% overlapped with transcripts detected at the mRNA level, demonstrating substantial concordance between the transcriptomic and proteomic datasets. However, this overlap was not preserved within statistically dysregulated molecules (p value < 0.05, unadjusted). Only five genes—Hspa5, S100a6, Plp1, Slc24a2, and Uggt1—were commonly dysregulated at both the mRNA and protein levels (Figure 4 A-B). In addition to the complexity of EVs, this limited concordance likely reflects the influence of posttranscriptional and posttranslational regulatory mechanisms, including mRNA stability, translational efficiency, and protein turnover, which are well documented contributors to the discordance between transcriptomic and proteomic data.

**Figure 4.**
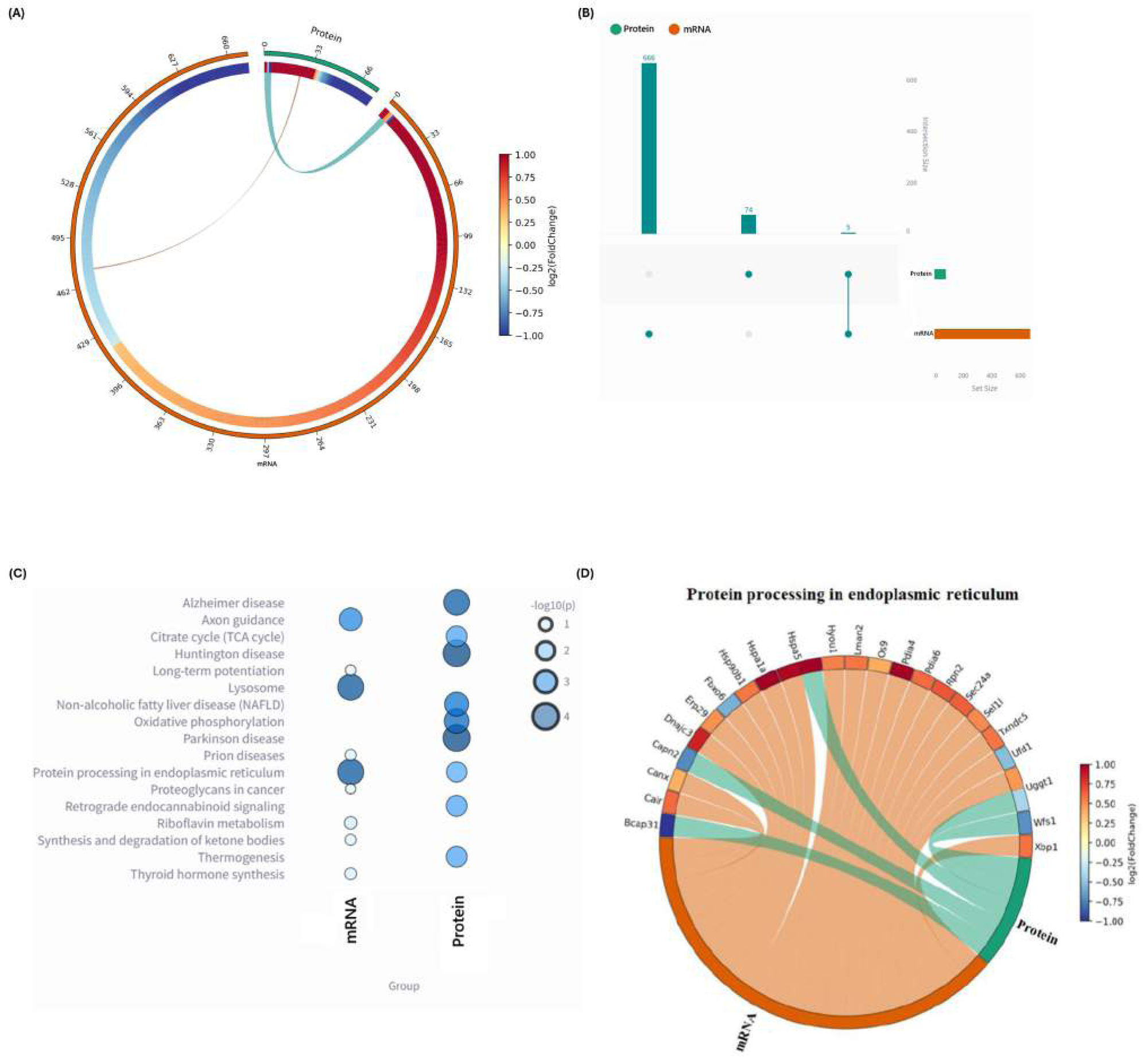
**Integrated transcriptomic and proteomic analysis of BDEVs from morphine-treated rats.** (A) Circos plot linking significantly dysregulated mRNAs (left) and proteins (right). Node colors represent log₂ fold-change (red/orange = upregulated, blue = downregulated). (B) UpSet plot showing the overlap between significantly altered RNAs and proteins; bar height represents the number of molecules in each set. (C) Comparative KEGG pathway enrichment analysis of dysregulated mRNAs and proteins. Bubble size reflects the number of molecules per pathway, and color indicates –log₁₀(p). (D) Chord diagram of the protein processing in endoplasmic reticulum pathway, depicting concordance and divergence between RNA and protein levels. Ribbons connect matched RNA–protein pairs, with ribbon color indicating log₂ fold-change.

Among the enriched pathways identified independently from the transcriptomic and proteomic datasets, protein processing in the endoplasmic reticulum was the only pathway common to both analyses. Interestingly, within this pathway, Hspa5 is upregulated at both the mRNA and protein levels in response to morphine exposure in BDEVs, suggesting conserved stress‒response activation [26,27]. In contrast, other transcripts contributing to this pathway were upregulated at the mRNA level, while their corresponding proteins were downregulated, further supporting the role of translational regulation or differential protein degradation in shaping the exosomal proteome under the influence of morphine (Figure 4 C-D) [28,29].

### Morphine-BDEVs modulate neurodegenerative gene expression and activate opioid signaling in cross-species recipient neurons

To investigate the functional role of BDEVs and their potential to modulate neurodegenerative processes, we treated primary neurons with morphine-BDEVs and control-BDEVs, followed by a comprehensive multiplex gene expression analysis targeting 770 neuropathology-associated genes **(Fig. 5A)**. After controlling for multiple comparisons, we identified significant differential expression of 15 genes in primary neuronal cells following treatment with morphine-derived BDEVs (FDR q value < 0.1). Among these genes, *Mnat1*, a gene implicated in blood‒brain barrier (BBB) damage in Alzheimer’s disease (AD), was downregulated, whereas *PSEN1*, *NSF*, and *NRG1*, which are associated with neurodegenerative processes, particularly in AD, were upregulated **(Fig. 5B)**. This evidence supports previous findings suggesting that extracellular vesicles (EVs) may contribute to the neurodegeneration and dementia observed with long-term opioid use [30].

**Figure 5.**
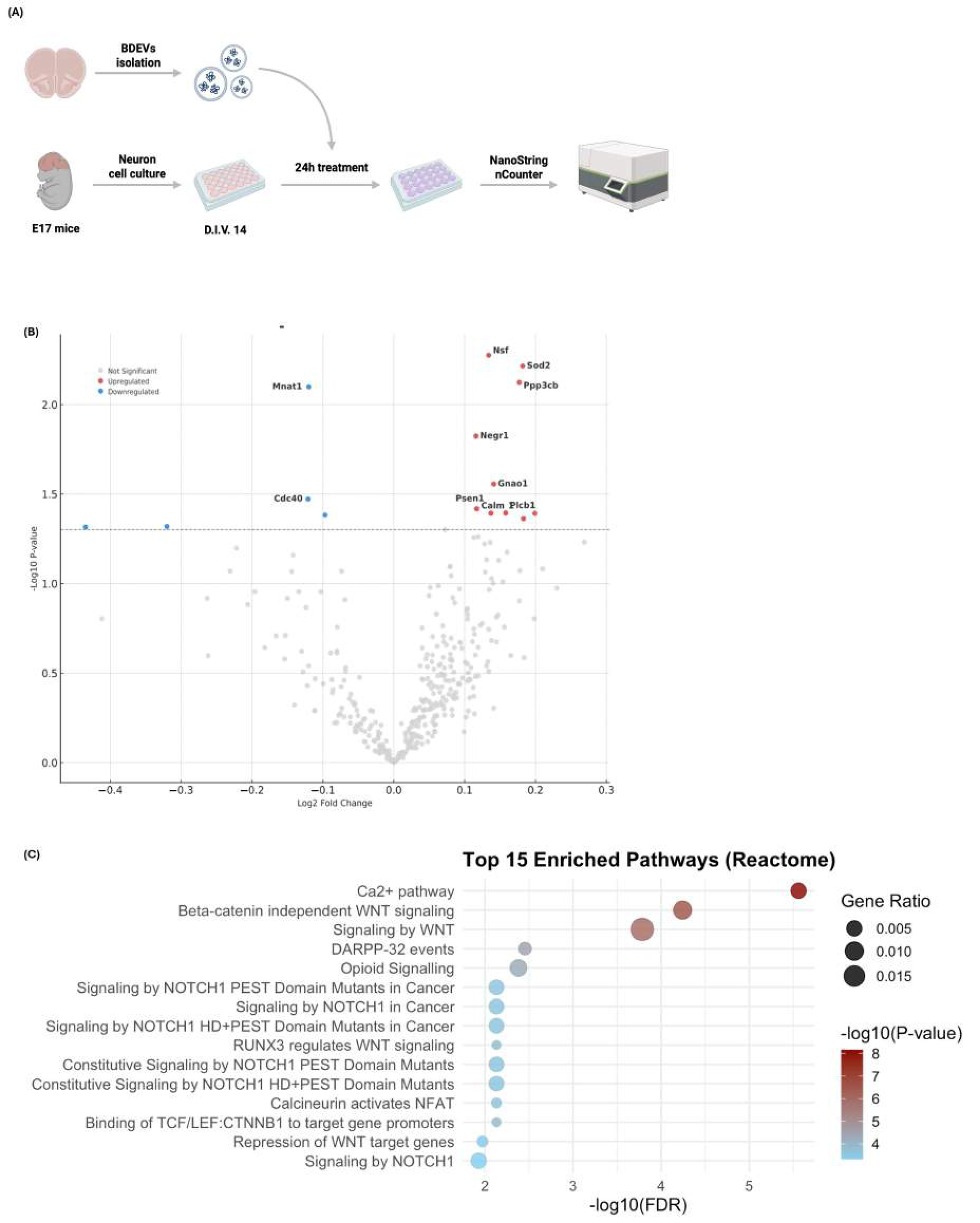
**Effect of morphine-derived BDEVs on primary neuronal cultures.** (A) Experimental design: BDEVs from morphine- and vehicle-treated rats were applied for 24 h to primary cortical neurons, followed by NanoString analysis. (B) Volcano plot showing differentially expressed transcripts in neurons exposed to morphine-derived BDEVs; selected significant genes are labeled. (C) Reactome pathway enrichment of dysregulated genes, highlighting Ca²⁺, WNT, opioid, and DARP 32-related signaling pathways.

Pathway analysis revealed a selective enrichment of non-canonical WNT signaling components in neurons exposed to BDEVs derived from morphine-treated rats, despite the absence of direct morphine exposure. The top-ranked pathway was “Ca^2+^ pathway” (FDR = 2.75 × 10⁻⁶; ratio = 0.005), followed by “Beta-catenin independent WNT signaling” (FDR = 5.71 × 10⁻⁵; ratio = 0.01), and “Signaling by WNT” (FDR = 1.64 × 10⁻⁴; ratio = 0.02). These non-canonical pathways are critical for cytoskeleton remodeling, intracellular calcium flux, and neuronal polarity. Remarkably, neurons also showed enrichment for “DARPP-32 events” and “Opioid signalling” (both FDR = 0.004), indicating activation of dopaminergic integration and GPCR-related neuromodulatory cascades. These results suggest that morphine-induced BDEVs can transfer biologically active signals capable of reprogramming recipient neurons toward a phenotype consistent with drug exposure, engaging both polarity-regulating pathways and classical neuroadaptive circuits **(Fig. 5C)**.

## Discussion

This study identifies BDEVs as active participants in the cascade of neuroadaptations induced by chronic opioid exposure. Using an integrative omics approach, we show that morphine reprograms the transcriptomic and proteomic cargo of BDEVs in the rat PFC, enriching them with molecules linked to synaptic plasticity, endoplasmic reticulum (ER) stress, mitochondrial dysfunction, and neurodegenerative processes. Importantly, functional assays demonstrated that BDEVs derived from morphine-treated animals were sufficient to alter transcriptional programs in naïve cortical neurons, including genes involved in excitability and structural remodelling. These findings advance our understanding of OUD pathophysiology by implicating vesicle-mediated intercellular communication as a novel mechanism of drug-induced plasticity.

### BDEVs as mediators of opioid-induced neuroadaptations

Addiction is widely recognized as a disorder of maladaptive plasticity, where repeated drug exposure induces long-lasting changes in brain circuits that regulate reward, cognition, and stress [31,32]. The PFC is a central hub for such adaptations, integrating glutamatergic and GABAergic inputs to modulate drug-seeking behaviour [33,34]. Our results demonstrate that chronic morphine alters the cargo of PFC-derived BDEVs, including transcripts and proteins central to synaptic remodelling (e.g., ARC, NCAM1), ER stress (Hspa5), and mitochondrial function (NDUF family) [35–37]. This suggests that opioids reshape not only intracellular signalling cascades but also vesicle-mediated communication between neural cell types, adding a new layer to established models of opioid-induced neuroplasticity.

The identification of ARC gene as a consistently upregulated molecule in BDEVs of animals treated with morphine is particularly significant. ARC is an important regulator of synaptic plasticity and has been linked to drug-induced synaptic remodelling in several brain regions, including PFC [36,38]. ARC has also been implicated in the biogenesis and release of EVs. It can self-assemble into mRNA-containing, virus-like capsids that are packaged into EVs and released by neurons, where they can be endocytosed by recipient cells at postsynaptic sites [13,39,40]. Thus, the selective enrichment of ARC in BDEVs suggests a mechanism by which morphine-exposed neurons may broadcast plasticity-related signals to neighbouring cells, thereby coordinating circuit-wide adaptations [40]. Similarly, upregulation of Hspa5 in vesicles aligns with previous studies showing opioid-induced ER stress in mesocorticolimbic structures, which has been implicated in tolerance and dependence [41–44]. Together, these findings support the notion that vesicles carry molecular cargo that reflects and potentially amplifies core processes driving OUD.

### Relationship to previous EV studies in opioid exposure

Previous research on EVs in the context of opioid exposure has primarily focused on peripheral vesicles, particularly blood-derived EVs enriched in microRNAs. For instance, opioid exposure has been associated with altered circulating miRNA profiles in both humans and animal models, while perinatal opioid exposure has been shown to modulate EV-associated small RNAs in the developing brain [45–47]. However, these studies were largely restricted to small RNA species, limiting insights into the broader transcript diversity of EVs, including messenger RNAs [48].

In contrast, our approach employed total RNA sequencing with ribosomal RNA removal, enabling the identification of mRNAs within BDEVs, consistent with recent evidence demonstrating the functional transfer of mRNAs via EVs [12], along with unbiased proteomic profiling. We further validated functional relevance by showing that morphine-altered BDEVs can reprogram transcriptional activity in recipient neurons. This extends beyond biomarker discovery, establishing BDEVs as active effectors of opioid-induced neuroadaptation.

Moreover, cross-referencing our RNA-seq data with a large-scale human GWAS meta-analysis for opioid use disorder revealed an overlap between morphine-altered vesicle cargo and genes associated with OUD, possibly underscoring the translational relevance of our findings [15]. Transcripts such as Oprm1, Ncam1, and Kcnn1 not only play established roles in synaptic function and neuronal excitability but also appear within genetic risk loci for OUD, suggesting that BDEV cargo reflects both drug-induced molecular changes and intrinsic susceptibility pathways [15,49–51].

### BDEV-mediated mechanisms of synaptic plasticity and circuit dysfunction in OUD

Our proteomic analyses further demonstrated significant alterations in the cargo of BDEVs following chronic morphine exposure, identifying dysregulated proteins implicated in neuroadaptations associated with opioid exposure. Specifically, we observed increased levels of PDIA3, STX1B, YWHAG, and HSPA5, molecules previously associated with synaptic dysfunction and pathways relevant to neurodegenerative diseases [52–55]. These findings align with previous evidence suggesting that prolonged opioid exposure may elevate the risk of neurodegeneration through mechanisms involving extracellular vesicles [56]. For instance, increased levels of neurodegeneration-related protein biomarkers have been detected in EVs isolated from the plasma of monkeys exposed to oxycodone [46]. Similarly, morphine-treated astrocytes have been shown to secrete EVs enriched with amyloid-related proteins, contributing to neurodegenerative processes in vitro [57].

Our pathway enrichment analyses across both proteomic and transcriptomic datasets revealed alterations in biological processes critical for synaptic remodelling, including axon guidance, lysosomal activity, and ER stress [58]. Alterations in axon guidance molecules have been reported in animal models of oxycodone self-administration, suggesting that vesicle-mediated signalling could contribute to aberrant synaptic connectivity and neuroplastic remodelling [59]. Likewise, the enrichment of ER stress pathways supports the hypothesis that opioid-induced plasticity involves maladaptive cellular stress responses [60].

Beyond ER stress, our analyses also revealed alterations in mitochondrial pathways, suggesting that morphine-evoked vesicle remodelling may broadly impact cellular energy metabolism. Notably, we observed enrichment of mitochondrial proteins such as COX1, COX7A2L, and NDUF subunits within BDEVs, suggesting that vesicle-mediated signalling may link opioid exposure to bioenergetic stress [61–63]. Mitochondrial dysfunction and oxidative damage are well-established consequences of chronic opioid use and have been implicated in both tolerance and neurotoxicity [22,64]. The selective export of mitochondrial proteins through BDEVs may reflect ongoing metabolic stress and could propagate vulnerability signals to neighbouring cells, thereby amplifying circuit dysfunction [65].

This interpretation is supported by our functional assays, which demonstrated that morphine-derived BDEVs were sufficient to induce transcriptional changes in naïve neurons consistent with neuroadaptive processes. A targeted gene expression panel revealed significant alterations in several neuropathology-associated genes, including upregulation of Psen1, Nsf, and Negr1, genes linked to neuronal connectivity, synaptic plasticity, and neurotransmitter signalling [66–68]. These findings indicate that morphine-BDEVs can trigger transcriptional programs aligned with neuroadaptive signatures. Pathway-level analysis further identified activation of opioid signalling and DARPP-32 cascades, as well as non-canonical WNT and calcium pathways associated with spine remodelling [69–71]. Collectively, these results suggest that BDEVs act as vehicles of synaptic reprogramming, transmitting drug-induced plasticity signals in the absence of direct receptor activation.

### Translational implications: BDEVs as biomarkers and therapeutic targets

The ability of BDEVs to encapsulate and transmit opioid-induced molecular signatures has important translational implications. First BDEV cargo may serve as a biomarker for OUD. Because EVs can cross the blood–brain barrier and be isolated from peripheral fluids, profiling vesicle cargo could provide a minimally invasive means to monitor brain changes associated with opioid exposure [72]. Second, vesicle-mediated communication may represent a novel therapeutic target. Strategies aimed at modulating EV release, uptake, or cargo loading have shown promise in other disease contexts [73–75]. If BDEVs are found to amplify or sustain maladaptive signaling in vivo, interventions that alter vesicle biology could mitigate drug-induced neuroadaptations [72]. For example, targeting ER stress pathways (e.g., Hspa5) or synaptic regulators (e.g., ARC) within vesicles may reduce the persistence of opioid-induced plasticity In conclusion, our findings identify BDEVs as active mediators of opioid-induced neuroplasticity. Chronic morphine exposure reprograms vesicle cargo at both transcriptomic and proteomic levels, enriching molecules involved in synaptic plasticity, stress responses, and mitochondrial function. These vesicles are not passive byproducts, but are capable of reprogramming recipient neurons, thereby contributing to the persistence of drug-induced adaptations. By linking BDEVs biology with addiction neuroscience, this work advances mechanistic understanding of OUD and highlights possible new opportunities for biomarker discovery and therapeutic intervention.

## Authors Contribution

PHG and RGO designed the study. PHG, SB, RR, FC, and NRS performed experiments and data acquisition. MRL and VCC conducted proteomic analyses. YY and TWV contributed to RNA sequencing and data processing. TB performed TB performed transmission electron microscopy. CBV, HKM, SJ, RGO provided supervision, resources, and critical input on experimental design and interpretation. RGO conceived the project, secured funding. PHG and RGO wrote the manuscript with input from all authors. All authors reviewed and approved the final version

