## Supplementary Materials for "Morphine reprograms brain-derived extracellular vesicles from cargo linked to synaptic remodelling in the prefrontal cortex" (from another version of this preprint: version 1, 2025-09-17)

**Full Methods**

**Animals**

Eight-week-old male Sprague–Dawley rats (RjHan:SD; JANVIER Labs, Le Genest-Saint-Isle, France) were habituated to housing conditions 2 weeks before testing. The rats were housed in pairs under standard laboratory conditions, with ad libitum access to food and water. This study complied with Directive 2010/63/EU and the Danish Act on Animal Experimentation (Consolidated Act No. 474 of 15 May 2014) and was approved by the Danish Animal Experiments Inspectorate (Dyreforsøgstilsynet; permit no. 2023-15-0201-01528).

**Drug treatment and tissue harvest**

Morphine hydrochloride was prepared in 0.9% sodium chloride. The repeated morphine administration paradigm was performed via a previously established procedure with minor modifications (1). Briefly, the rats were injected intraperitoneally (i.p.) with saline or an increasing dose of morphine (from 10 mg/kg to 50 mg/kg) twice daily for 5 consecutive days (**Fig. 1 A**). Thus, the total dose of morphine used was 20–100 mg/kg daily (10 injections total). Four hours after the animals received their final dose of morphine, they were sacrificed by decaptation. The brains were removed, and the bilateral prefrontal cortex(PFC) was dissected and quickly frozen on dry ice, and stored at −80 °C until use.

**Brain-derived extracellular vesicle isolation**

BDEVs were isolated as previously described (2). Briefly, PFCs samples (∼90 mg) were minced and incubated with 20 U/mL papain (Worthington, Lakewood, NJ, US) in Hibernate A (BrainBits, Springfield, IL, US) for 15 min at 37 °C. The enzymatic digestion was stopped by the addition of ice-cold protease inhibitors to Hibernate A. The mixture was pipetted to further loosen the tissue and then centrifuged at 300 × g for 10 minutes at 4 °C. The supernatant was filtered sequentially, first, using a 40 μm cell strainer, followed by filtration through a 0.2 μm surfactant-free cellulose acetate (SFCA) syringe filter. Then, we centrifuged the solution obtained from filtration at 2,000 × g for 10 min at 4 °C, and the supernatant was centrifuged at 10,000 × g for 30 min at 4 °C. The supernatant was then ultracentrifuged at 100,000 × g (k factor: 207.5, 45Ti rotor type, Beckman Coulter, Brea, CA, US) for 70 min at 4 °C. The pellet was washed once in PBS, pH 7.4 (Corning Inc.), and recentrifuged at 100,000 × g for 70 min at 4 °C. The resulting crude EV pellet was resuspended in 500 µL of PBS. To characterize the extracted BDEVs, a BCA protein assay, Western blot, transmission electron microscopy (TEM), and nanoparticle tracking analysis (NTA) were performed, as detailed in the following sections.

**Transmission electron microscopy (TEM)**

Electron microscopy was performed as previously described (3). Briefly, three microliters of a diluted BDEV sample was applied to a 400-mesh collodion-coated (Sigma‒Aldrich) and carbon-coated copper grid (Pelco), which had been glow-discharged for 45 s at 25 mA and 39 mbar using an EasiGlow device (Pelco). After 30 s of incubation, the grid was blotted with 85 mm Grade 1 filter paper and subjected to three cycles of washing and staining with 3 µL of 2% uranyl formate (Polysciences Europe GmbH), with blotting steps between each washing/staining step and a final blotting and drying step. Micrographs were acquired via a Tecnai Spirit TWIN transmission electron microscope (Thermo Fisher Scientific) operated at 120 kV. Images were captured with a TemCam F416 CMOS camera via EM-Menu software (Tvips).

**Nanoparticle tracking analysis (NTA)**

The particle concentration and size distribution were determined via NTA via an LM14 NanoSight instrument (Malvern Instruments, UK). The samples (diluted 1:1000) were analyzed in triplicate (405 nm, camera level 16, syringe pump speed 20) and processed via Nanosight NTA software v. 3.0 (Malvern Pananalytical).

**Western blot analysis**

The purity of the isolated BDEVs was assessed via western blotting with specific antibodies as described previously (4, 5). Briefly, exosome protein lysates were prepared in RIPA buffer supplemented with 1% SDS and a protease-phosphatase inhibitor. Protein quantification was carried out via the Pierce BCA Protein Kit (Thermo Scientific) according to the manufacturer’s guidelines. Next, 25 microlitres of whole-brain and BDEV samples were loaded into an SDS‒polyacrylamide gel (26 wells—NuPAGE 10% Bis‒Tris Midi Gel—Thermo Scientific). The separated proteins were transferred to a 0.2 µM nitrocellulose membrane (Midi format, Bio-Rad) via the Bio-Rad Trans-Blot Turbo Transfer System. The membrane was blocked in intercept blocking buffer (IBB; LICOR) for one hour at room temperature and then incubated with a primary antibody diluted in IBB/0.1% Tween-20 at 4 °C overnight: with primary antibodies against the positive BDEV markers Hsp70 (1:1000, Sigma), flotillin-1 (1:1000, Abcam), and the negative marker GM-130 (1:1000, Abcam), followed by a secondary antibody (1:2500 HRP-conjugated anti-mouse IgG for Hsp70, 1:2500 HRP-conjugated anti-rabbit IgG for flotillin-1 and GM-130). The membranes were washed four times in TBS with 0.1% Tween-20 and once with TBS buffer for 5 min. The protein levels were assessed via Odyssey CLx Scan (LICOR).

**BDEV RNA purification and genomic DNA removal**

We used 20 µl of BDEVs for RNA purification via the miRNeasy Serum/Plasma Advanced Kit (QIAGEN) according to the manufacturer’s protocol, and the RNA was eluted in 50 μl of RNase-free water. The genomic DNA was removed via the Turbo DNA-free Kit (Ambion, MA, USA) following the manufacturer’s instructions. From the treated sample, 50 μl of DNA-free RNA was obtained. To concentrate the RNA, 50 μl of RNase-free water was added, followed by the addition of 10 µl of 3 M sodium acetate (pH 5.5), 250 µl of prechilled 99% ethanol, and 1 µl of GlycoBlue (Ambion, MA, USA). The mixture was incubated at −20 °C overnight. The RNA was then pelleted by centrifugation at 16,000 × g for 20 minutes at 4 °C. The resulting pellet was washed with 1 ml of 80% ethanol. Following a second centrifugation at 10,000 × g for 5 minutes, the RNA pellet was resuspended in 12 µl of RNase-free water. The RNA quality and quantity were assessed via a Bioanalyzer RNA 6000 Pico Kit (Agilent). The RNA was subsequently stored at −80 °C.

**BDEV total RNA sequencing**

The total RNA sequencing library was constructed via the use of 2 ng of purified RNA, following the SMARTer Stranded Total RNA-Seq Kit v3—Pico Input Mammalian protocol (Takara, Japan). Fragmentation was carried out for 90 seconds, with the remainder of the experiment conducted according to the instructions. The size distribution of the library fragments was assessed via the Bioanalyzer High Sensitivity DNA Analysis Kit (Agilent), and the library concentration was quantified via the Qubit dsDNA HS (High Sensitivity) Assay Kit (Thermo Fisher). The libraries were then pooled in equal amounts and sequenced on a NovaSeq X Plus sequencing platform via a 10B flow cell for paired-end 150-cycle sequencing (Illumina, MA, USA).

**RNA sequencing data analysis**

To estimate cell-type proportions in each RNA-seq sample, we employed the CIBERSORT R package. The reference dataset for this analysis was derived from the Allen Brain Map Mouse Cortex and Hippocampus SMART-seq single-cell RNA sequencing data, which included profiles for astrocytes, microglia, oligodendrocytes, glutamatergic pyramidal (PT) neurons, glutamatergic intratelencephalic (IT) neurons, glutamatergic corticothalamic (CT) neurons, glutamatergic near-projecting (NP) neurons, and GABAergic interneurons. We then used all the RNA-seq counts from each sample as input. RNA-seq targets with a base mean value less than 5 were excluded from the analysis because of insufficient counts. Differential gene expression (DGE) analysis was conducted via DESeq2 (version 1.34.0) (6). Dispersion estimates were obtained through a negative binomial model, followed by Wald tests to evaluate differential expression. The significance thresholds for DGE were determined via Benjamini–Hochberg false discovery rate (FDR) correction, with an adjusted p value < 0.1 and a log2-fold change (log2FC) threshold of ±0.5. Gene Ontology (GO) enrichment analysis was conducted on the list of genes with unadjusted p values < 0.05 via the Metascape platform (Metascape.org). This analysis focused specifically on terms related to the Cellular Component and Biological Process categories. Statistical significance thresholds for GO analysis were set at a p value < 0.01, with a requirement of at least 3 overlapping genes/proteins between the input list and the pathway list.

**Proteomic analysis of BDEVs**

BDEVs from control and morphine-exposed rats were lysed in HEPES buffer (pH 8.0) supplemented with 2% sodium deoxycholate (SDC). The protein concentration was measured via a Nanodrop, and buffer contaminants were removed via C8 stage tips. Proteins were reduced with DTT, alkylated with IAA, and digested overnight with trypsin. The peptides were desalted on C18 microcolumns and resuspended in 0.1% formic acid. For liquid chromatography‒tandem mass spectrometry (LC‒MS/MS), 0.5 µg of peptide was separated on a C18 analytical column via a 90-min gradient (3–40% ACN). An Orbitrap Q Exactive HF operated in DDA mode with 120K resolution for MS1 and 15K resolution for MS2, selecting the 15 most intense ions for HCD fragmentation. The raw data were searched against the *Rattus norvegicus* UniProt database via Proteome Discoverer (MASCOT) with a 1% FDR. Proteins were quantified on the basis of at least one unique peptide and ≥3 peptides across 80% of the samples. For protein interaction and pathway enrichment analysis, Cytoscape and R-based in silico analyses were used. Principal component analysis (PCA) and volcano plot analysis were conducted via OmicScope (7). Ingenuity pathway analysis (IPA) was used for protein function integration, with a significant cutoff of a fold change (FC) >2.0 or <-2.0 and p < 0.05.

**Primary Neuronal Culture**

Cell cultures were prepared from embryonic day 17 (E17) of C57BL/6 embryos, following euthanasia of pregnant females under approved ethical protocols, as previously described (8). Briefly, 24-well plates were precoated with 0.5 mg/mL poly-L-lysine (Sigma) prepared in B buffer (3.1 mg/mL boric acid and 4.75 mg/mL borax; Sigma) for 4 hours at 37 °C with 5% CO₂. Plates were then washed with sterile deionized water and PBS and covered with plating medium. The neocortices from E17 embryos were pooled, and minced before enzymatic digestion in 2.5% trypsin and DNase I at 37 °C for 15 minutes. Next, the tissue was washed with neuroplating medium and mechanically dissociated using a Pasteur pipette for 10-15 strokes and a fire-polished Pasteur pipettes for additional 10-15 strokes until large clumps were reduced, and the suspension was allowed to settle. The upper cell suspension was collected, and the trituration process was repeated twice on the remaining clumps. The combined cell suspension was filtered through a 70 μm cell strainer and the cells were counted by Luna-II automated cell counter (Logos Biosystems). To separate glial cells from neurons, the suspension was incubated in an uncoated Petri dish at 37 °C with 5% CO₂ for 30–60 minutes, allowing glial cells to preferentially adhere. The neuronal cell suspension was centrifuged, and the cells were resuspended in Neurobasal medium supplemented with B27. Neurons were plated at a density of 5 × 10⁵ cells/mL. Half of the medium was replaced every third day. Neuronal culture experiments were performed on day in vitro (DIV) 7.

**NanoString nCounter® Gene Expression**

Primary cortical neurons were treated with 1 × 10⁸ BDEVs isolated from the prefrontal cortex of morphine- or vehicle-treated rats. BDEVs were resuspended in sterile PBS and added directly to the culture medium. Twenty-four hours after treatment, cells were collected and processed for transcriptomic profiling using the NanoString nCounter® system. Total RNA was isolated using the miRNeasy Mini Kit (Qiagen, Germany) according to the manufacturer’s instructions. RNA quantity and purity were assessed with a NanoDrop-1000 spectrophotometer (Thermo Scientific, USA). Gene expression was quantified using the NanoString nCounter® Mouse Neuropathology Panel (XT-CSO-MNROP1-12; Table S3), which targets 770 genes associated with neurodegeneration, following the manufacturer’s protocol. For each sample, 30 ng of RNA was hybridized with the codeset at 65 °C for 16 h. After automated processing using the nCounter® Prep Station, data acquisition was performed on the nCounter® Digital Analyzer. Gene expression was normalized to internal housekeeping genes and analyzed using nSolver™ Analysis Software 4.0 (NanoString Technologies). Six independent samples per group were used (primary neuronal cultures treated with morphine-BDEVs vs. saline-BDEVs). Following NanoString guidelines and previously validated methodology (Veldman-Jones et al., 2015), differential expression was assessed using nominal p-values, without correction for multiple comparisons, given the targeted nature of the panel and its predefined biological relevance. All genes with nominal p-values < 0.05 (n = 22) were selected for pathway enrichment analysis using the Reactome Pathway Database (version 88). Over-representation analysis was conducted via Reactome’s online platform, using Homo sapiens as reference species. The statistical significance of pathway enrichment was determined by a binomial test with Benjamini–Hochberg correction for multiple comparisons. Enriched pathways were ranked by FDR, and visualized as dot plots with gene ratio and –log10(p-value) metrics.

**Cell viability assay**

***Methods***
To assess the cytotoxicity of varying concentrations of brain-derived extracellular vesicles (BDEVs), we used the CellTiter-Glo® 2.0 Luminescent Cell Viability Assay (Promega) following the manufacturer’s protocol(reference). Primary neuronal cells from E17 mice (collected as described in the main Methods) were treated for 24 h with: Vehicle (PBS), BDEVs isolated from the prefrontal cortex of saline-treated or morphine-treated rats, at 10⁷, 10⁸, or 10⁹ particles/mL and 20% DMSO (positive control for cell death). Luminescence was measured on a Varioskan microplate reader and quantified using SkanIt software.

**Results**
As expected, treatment with 20% DMSO produced a drastic loss of viability in primary neurons (F(8, 54) = 2,632; p<0,0001; Supplementary Figure 1). In contrast, all concentrations of BDEVs, regardless of whether they were derived from saline- or morphine-treated animals had no significant effect on cell survival relative to vehicle (PBS).

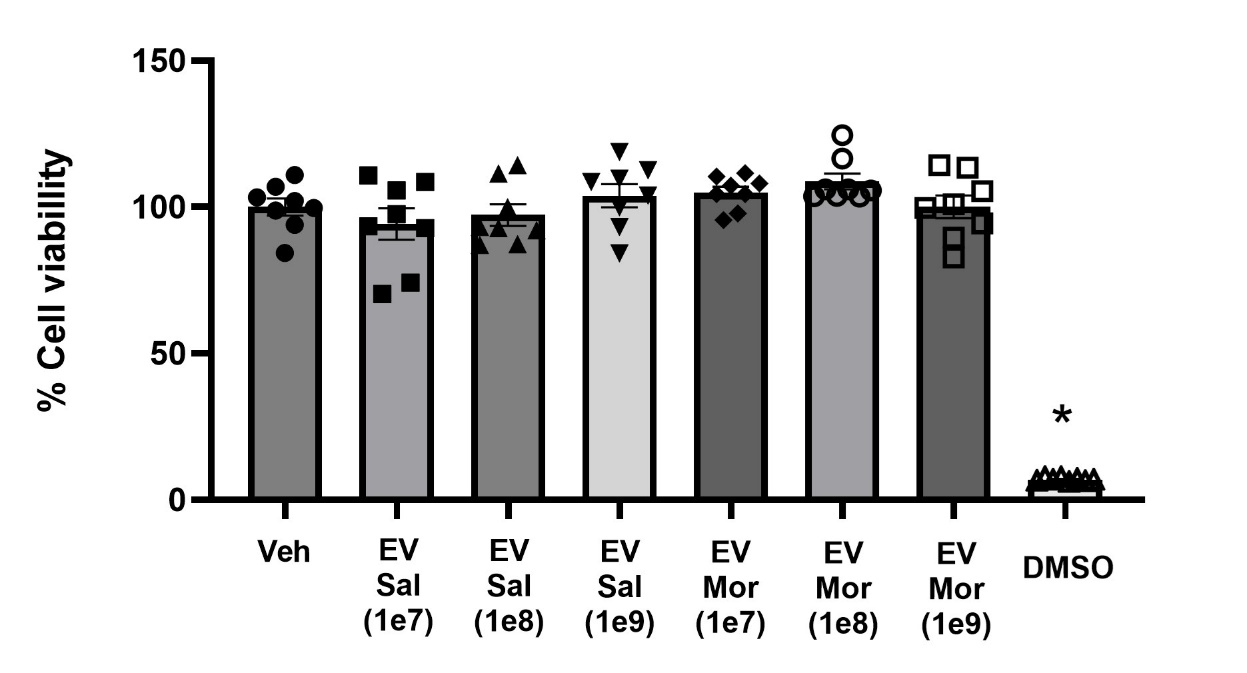

**Suplementar Figure 1**. Cell viability of primary neuronal cells from E17 mice treated with different BDEVs from morphine or saline in crescent concentrations (24h treatment). Significant decrease in neuronal viability was observed in the group of cells treated with DMSO (p < 0.0001, one-way ANOVA followed by Dunnett’s post hoc, n = 8)

8.	Sahu MP, Nikkila O, Lagas S, Kolehmainen S, Castren E (2019): Culturing primary neurons from rat hippocampus and cortex. *Neuronal Signal*. 3:NS20180207.
